# Hybrid Modeling of Engineered Biological Systems through Coupling Data-Driven Calibration of Kinetic Parameters with Mechanistic Prediction of System Performance

**DOI:** 10.1101/2023.06.14.545039

**Authors:** Zhang Cheng, Avner Ronen, Heyang Yuan

## Abstract

Mechanistic models can provide predictive insight into the design and optimization of engineered biological systems, but the kinetic parameters in the models need to be frequently calibrated and uniquely identified. This limitation can be addressed by integrating mechanistic models with data-driven approaches, a strategy known as hybrid modeling. Herein, we developed a hybrid modeling strategy using bioelectrochemical systems as a platform system. The data-driven component of the model consisted of artificial neural networks (ANNs) that were trained by using mechanistically derived parameter values (e.g., the maximum specific growth rate µ_max_ and the maximum substrate utilization rate k_max_ for the fermentative, electroactive, and methanogenic populations, and the mediator yield for electroactive microbes Y_M_) as outputs to compute error signals. The hybrid model was built using 148 samples collected from 25 publications. After ten-fold cross-validation, the model was tested with another 28 samples. Internal resistance was accurately predicted with a relative root-mean-square error (RMSE) of 3.9%. Microbial kinetic parameters were also calibrated using the data-driven component. They were fed into the mechanistic component to predict system performance. The R^2^ between the predicted and observed organic removal and current production for systems fed with a simple substrate were 0.90 and 0.94, respectively, significantly higher than those obtained with a standalone data-driven model (0.51 and 0) and a standalone mechanistic model (0.07 and 0.15). The hybrid modeling strategy can potentially be applied to a variety of engineered biological systems for *in silico* system design and optimization.

**SYNOPSIS:** A hybrid modeling strategy was developed to predict the performance of engineered biological systems without the need for laborious experiment-based parameter calibration.

## 1. INTRODUCTION

Engineered biological systems, such as activated sludge processes and anaerobic digesters, have been widely used to treat waste streams and recover valuable resources,^1^ providing sustainable solutions to the challenges at the water-energy nexus.^2-4^ There is also continued interest in developing new biotechnologies, e.g., bioelectrochemical systems,^5-7^ to achieve more efficient treatment and recovery with lower energy consumption and carbon emissions. The design and operation of those systems remain challenging because the microbial processes involved are highly sensitive to environmental and operational variations, such as the changes in substrate composition and concentration, electrode material and catalyst, operating mode, external resistance, reactor configuration, and temperature, etc.^8, 9^ The main influencing factors vary distinctly for bioelectrochemical systems with different configurations, operating modes, and applications. For example, the internal resistance of a microbial desalination cell can change from tens to hundreds of ohms due to the decrease in salinity in the desalination chamber.^10-12^ A saline anode can promote electroactivity, but excess salinity may inhibit microbial growth.^13, 14^ This has motivated the development of mechanistic models for predictive control of engineered biological systems.

Mechanistic models (also known as first-principle or white-box models) are powerful tools for understanding, predicting, and optimizing system performance.^15-17^ Built based on the mathematical descriptions of the fundamental principles of the systems, mechanistic models can generate interpretable predictions for exploring the underlying mechanisms that govern the behavior of engineered biological systems. Mechanistic models can also be used to identify key environmental conditions and test hypotheses, enabling the optimization of existing systems and the design of new systems. With over 60 years of development,^18, 19^ various mechanistic models have been built to model different engineered biological systems. Between 1987 and 1999, a task group from the International Association on Water Quality developed the standard mechanistic models ASM1/2/2d/3 to simulate the microbial processes in activated sludge systems.^20–23^ The standard mechanistic model ADM1, developed in 2002 by another task group, was able to describe the 19 microbial processes in anaerobic digesters.^24^ The same modeling strategy has also been used to model emerging biotechnologies such as membrane bioreactors, anaerobic ammonium oxidation processes, and bioelectrochemical systems.^15, 25–30^ For example, a model combining mass balance and the Nernst’s equation was proposed to predict the performance of anodic biofilm formed in microbial fuel cells. The model was constructed using one-dimensional differential equations to simulate substrate utilization, biomass inactivation, and respiration.^31^ In addition, the mass balances for electron transfer mediators and salts were added to develop a new mechanistic model for microbial desalination cells.^28^

Despite their wide implementation, mechanistic models suffer from laborious calibration and identification of kinetic parameters. Many of the parameters, in particular those related to microbial growth kinetics, are not measurable and need to be frequently calibrated with experimental data.^32^ Parameter calibration is typically performed using a trial-and-error approach, and the difference between the predicted and observed values is evaluated manually.^33, 34^ This conventional method is time-consuming and may not be reproducible. Parameter calibration can also be performed using automated algorithms,^35^ but the selection of the final parameter values relies heavily on expert knowledge. Calibration without expert supervision can result in inaccurate and uninterpretable predictions.^36^ Regardless of the calibration method, a unique set corresponding conditions, thus avoiding the need to identify a unique set of parameter value across conditions.

This promising strategy has not been implemented in practical applications possibly due to several limitations.^54–57^ First, data-driven approaches have been criticized for yielding uninterpretable inferences because of their poor extrapolation performance and black-box nature.^58^ This is particularly problematic when the information about microbial processes is not incorporated into model training.^59–63^ For example, the parameter values are sometimes inferred to be negative and must be manually set to zero.^56^ With such biologically unreasonabl parameter values, the downstream mechanistic models cannot yield accurate predictions or provide mechanistic insight. In addition, both environmental conditions and prior system outputs are time-independent variables that cannot adequately reflect time-dependent parameter values. As a result, the timeseries predictions from the hybrid model were less accurate than those from standalone data-driven models.^57^ Last but not least, the resulting hybrid models still rely on experimental data (i.e., prior system outputs) for parameter calibration.

We recently modified the hybrid modeling strategy to improve the interpretability of parameter inference.^64^ For the first time, microbial population abundances were included as the inputs. Another major change was to incorporate mechanistically derived parameter values into the training of the data-driven component. Additionally, Bayesian networks were used to form the data-driven component to infer the causal relationship between microbial population abundances and microbial kinetic parameters.^65^ The modified strategy was examined using 77 samples collected from various bioelectrochemical systems. After model construction, the data-driven component was able to infer the connection between well-characterized electroactive populations and current production-related parameters. Meanwhile, the mechanistic component could use the predicted parameters to achieve robust simulation of system performance over a wide range of conditions. However, similar to that in the original strategy, our data-driven component did not consider the time interval between the input and output and could not predict the dynamic behavior of microbial processes. Our strategy also required substantial experimental data as the input, in particular microbial population abundances which were often unavailable, limiting the applicability of the resulting model.

The goal of this work is to address the remaining limitations of the hybrid modeling strategy. To this end, the strategy was modified as follows (Figure 1B): 1) Artificial neural networks (ANNs) were used to form the data-driven component. Compared to Bayesian networks, ANNs can yield more accurate predictions due to their flexibility in dealing with high-dimensional inputs and nonlinear input-output relationships.^53, 66^ 2) Mechanistically derived parameter values were used as outputs to compute the error signals that were backpropagated for network training. This was expected to impose constraints on the calibrated parameter values and thus avoid uninterpretable inferences. 3) Environmental conditions were used as inputs, while all experimental data (e.g., prior system outputs and microbial population abundances) were excluded. 4) The time period over which parameter values were derived was included as an input. This time term acts as a filter that allows us to infer parameter values at different time points. Similar to our previous study,^64^ bioelectrochemical systems were used here as a platform system for examining the proposed strategy. Bioelectrochemical systems mainly consist of anode and cathode.^6, 67^ The electroactive microbes in the anode catalyze the conversion of organic matter into extracellular electrons.^68–70^ The electrons are transferred to the cathode for the reduction of a thermodynamically favorable electron acceptor (e.g., oxygen), resulting in current production. Internal resistance is a key parameter that is related to electron transfer kinetics and can strongly affect the dynamic behavior of bioelectrochemical systems.^5^ We also collected another 28 samples to test the robustness of the resulting model. Developed based on bioelectrochemical systems, the hybrid modeling strategy is expected to be broadly applicable to a variety of engineered biological systems.

**Figure 1.**
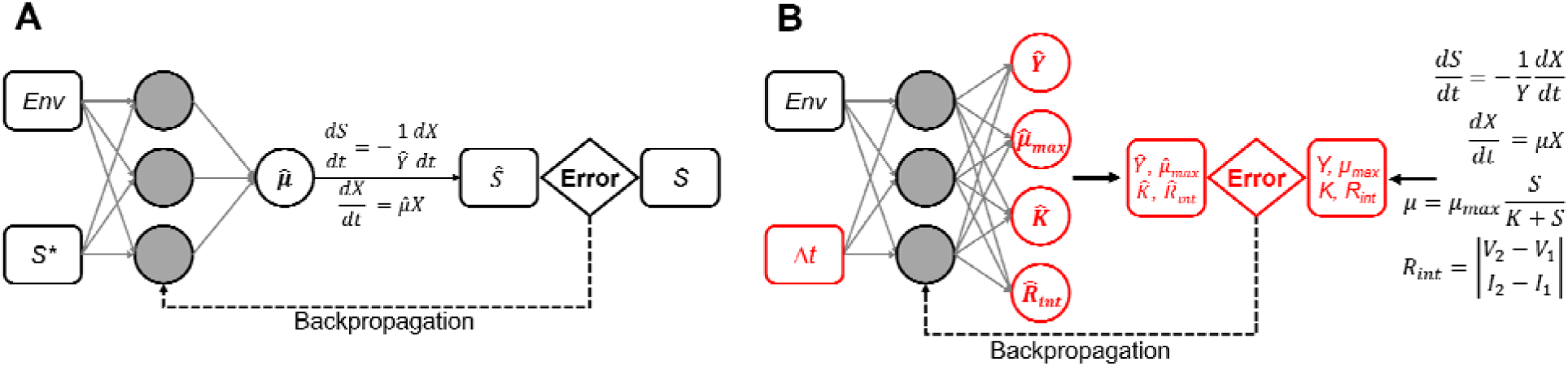
(A) Schematic of the hybrid modeling strategy proposed in an early study.^52^ (B) Schematic of the hybrid modeling strategy proposed in this study. The differences between the two strategies are highlighted in red. *Env*: environmental conditions; *S\**: prior system outputs; *S*: present system performance; Δ*t*: time interval; *µ*: growth rate; *µ_max_*: maximum specific growth rate; *K*: Monod constant; *Y*: yield; *R_int_*: internal resistance. The sign indicates inference.

## 2. METHODS

### 2.1 Collection of training dataset

We reviewed in total 169 publications and collected 148 samples from 25 publications for training the data-driven component [Supporting Information (SI) Table S1]. Because some of the samples lack key features to train the data-driven component for calibrating microbial kinetic parameters or internal resistance, the samples were divided into two groups to leverage the available data (some of the samples were used in both groups). Briefly, 73 samples from 12 publications were selected to train the ANNs for calibrating microbial kinetic parameters, and 86 samples from 17 publications were selected to train the ANN for calibrating internal resistance. To demonstrate the diversity of the samples, principal coordinate analysis (PCoA) based on Bray-Curtis distance was performed using the software R. More details about the collections of the training dataset can be found in SI Methods and Tables S2 & S3.

### 2.2 Derivation of kinetic parameter values

After collecting the training dataset, the kinetic parameter values were calculated in preparation for training the data-driven component. Real-time COD or current production was acquired directly from the figures in the publications using GetData Graph Digitizer 2.25.^71^ Microbial kinetic parameters were then determined using an established mechanistic model as described in our previous study.^64^ These include the maximum specific growth rate *µ_max_* and maximum substrate utilization rate k_max_ for the fermentative, electroactive, and methanogenic populations, and the mediator yield for electroactive microbes Y_M_. Internal resistance as a key parameter for electron transfer kinetics was calculated based on the polarization curves reported in the publications.^72, 73^ A polarization curve can be divided into three regions that represent activation loss, ohmic loss and concentration loss.^74^ The ohmic loss region, which is typically the middle part of a polarization curve, commonly shows a linear relationship between the voltage and current. The internal resistance can thus be estimated from the slope of the ohmic loss region. More details can be found in the SI Methods Internal resistance calculation.

### 2.3 Construction and validation of the data-driven component

Before the data-driven component was trained, all the variable values were scaled to between 0 and 1.^75^ Normalization is widely applied in data-driven modeling to make data of different scales comparable and to avoid bias.^76^ After normalization, the ANNs for calibrating microbial kinetic parameters were trained with operating conditions and reactor configuration as the inputs and mechanistically derived kinetic parameter values as the outputs (SI Table S2). The ANN for predicting internal resistance was trained with operating conditions, reactor configuration, and electrochemical conditions as the inputs and the calculated internal resistance as the output (SI Table S3). A time term (Δt in Figure 1B) was included as an additional training input based on the fact that the values of both microbial kinetic parameters and internal resistance could vary as a function of time. For microbial kinetic parameters, Δt is the hydraulic retention time. For internal resistance, Δt is the time period from the start of system operation to the measurement of internal resistance. For example, if a polarization curve was measured at the end of the operation cycle, Δt was then equal to the hydraulic retention time. If a polarization curve was measured immediately after a fresh substrate was added, Δt = 0. ANNs were trained using the R package Neuralnet.^77^ For simple training datasets, a three-layer ANN is sufficient to learn the statistical connection among the features. Empirically, the number of nodes in each hidden layer should be between the number of inputs and outputs.^78, 79^ Excessive hidden layers or nodes can lead to overfitting and compromise the prediction accuracy. Thus, the ANNs contained three hidden layers and up to eight nodes in each hidden layer. Considering the dataset size and computational cost, ten-fold cross-validation was used to validate the ANN training method.^80, 81^ Relative root-mean-square error (RMSE) and coefficient of determination (R^2^) were calculated as validation metrics. Null models were constructed as an additional validation step by averaging all the inputs, kinetic parameters, and internal resistance across all samples.^82^ More details about the training and validation procedures can be found in the SI Methods.

### 2.4 Prediction of system performance

To test the prediction performance of the hybrid model, 28 samples were collected from seven additional publications. Three types of system performance were predicted: internal resistance, COD removal, and current production. Internal resistance was predicted directly from the trained ANN and compared with the values derived from polarization curves. The time term (Δ*t* in Figure 1B) had to be determined based on the operation duration before the polarization curves were measured. For example, if a polarization curve was measured immediately after a fresh substrate was added, Δ*t* = 0. If a polarization curve was measured at the end of the operation cycle, Δ*t* was then equal to the hydraulic retention time. Microbial kinetic parameters were inferred with the time term determined using the same method. COD removal was then calculated by inputting the calibrated internal resistance and microbial kinetic parameters into the mechanistic component. The initial values for mechanistic prediction were assigned according to the method reported in the previous study.^64^ Current production was predicted in a dynamic manner using Ohm’s law. To achieve dynamic prediction, an operation cycle was manually divided into several intervals, and the internal resistance within each interval was inferred by feeding the ANN with the corresponding Δ*t*. Current production was therefore constantly updated within an operation cycle. Standalone ANNs were trained with the same inputs but COD removal and current production as the outputs. Standalone mechanistic prediction was also performed by feeding the established mechanistic model with averaged microbial kinetic parameters and internal resistance of the 141 samples. More details about the prediction procedures can be found in the SI Methods.

### 2.5 *In silico* design of bioelectrochemical systems

To further demonstrate the applicability of the hybrid model, a case study was conducted. The goal was to design a single-chamber microbial fuel cell to treat synthetic wastewater containing 600 mg/L acetate. Most of the features were set to typical values. These include room temperature (25 L), standard pressure (1 atm), fixed external resistance (10 Ω), etc. (SI Table S4). Five configuration-related features (anode/cathode area, cathode electrode conductivity, electrolyte conductivity, and electrode distance) and one operation-related feature (retention time) were considered to be key parameters and varied within certain ranges (SI Table S4). The ranges were determined based on the 169 publications reviewed in this study. The number of values of the five configuration-related features ranged from five to eight, resulting in a total of 11,120 combinations. Each combination was fed as the inputs into the ANN for internal resistance prediction, and the simulation was repeated 11,120 times. The combination that resulted in the minimum internal resistance was then fed along with retention time into the ANNs for calibrating microbial kinetic parameters. Six retention times ranging from 6 h to 192 h were used for the simulation (SI Table S4). Finally, the predicted internal resistance and kinetic parameters were input to the mechanistic component to calculate COD removal.

## 3. RESULTS AND DISCUSSION

### 3.1 Evaluating the sample diversity

The bioelectrochemical systems selected for examining the hybrid modeling strategy are highly diverse in terms of configuration and operation. In the training dataset, 93 samples were from microbial fuel cells, 38 were from microbial electrolysis cells, and 17 were from microbial desalination cells. Similarly, 23, 1, and 4 of the testing samples were collected from microbial fuel cells, microbial electrolysis cells, and microbial desalination cells, respectively. These bioelectrochemical systems were operated under a variety of conditions. For example, microbial fuel cells were fed with different substrates (e.g., simple fatty acids, fermentable sugars, or real wastewater) and operated under different modes (e.g., batch vs. continuous) with the external resistance ranging from 1 to 10,000 Ω. Microbial electrolysis cells were operated with the applied voltage ranging from 0.4 to 1.0 V under acidic, neutral, or alkaline conditions. For microbial desalination cells, the salinity in the desalination chamber differed by two orders of magnitude.

To quantitatively demonstrate the diversity of the selected systems, PCoA was performed with the training dataset for calibrating microbial kinetic parameters. This dataset consisted of operation-related features such as substrate composition and concentration and configuration-related features such as electrode distance and anode area (SI Table S2). As shown in Figure 2A, the bioelectrochemical systems fed with different substrates (i.e., non-fermentable and fermentable) and operated under different modes (i.e., batch, fed-batch, and continuous) were separated from each other. For example, the samples from Study no. 54 (S54) and S37 were fed with acetate and clustered on the right side in Figure 2A. In comparison, the samples from S48 and S20 were fed with ethanol and undefined complex organics, resulting in clear separation on the top and bottom left, respectively. PCoA was also performed with the training dataset for calibrating internal resistance (Figure 2B). This dataset contained distinct features such as electrode characteristics and ion strength (SI Table S3). The samples from S54 and S82 were separated from other samples because they were microbial desalination cells that had non-zero values of ion strength in the desalination chamber. The samples operated with high external resistance (e.g., S22, S51, S80, and S81) clustered at the bottom left in Figure 2B. In both PCoA graphs, the testing dataset scattered within and outside of the region of the training dataset, indicating its representativeness. Overall, the results show the high diversity of the selected bioelectrochemical systems and operating conditions, which is desirable for building a robust data-driven component in the hybrid model.

**Figure 2.**
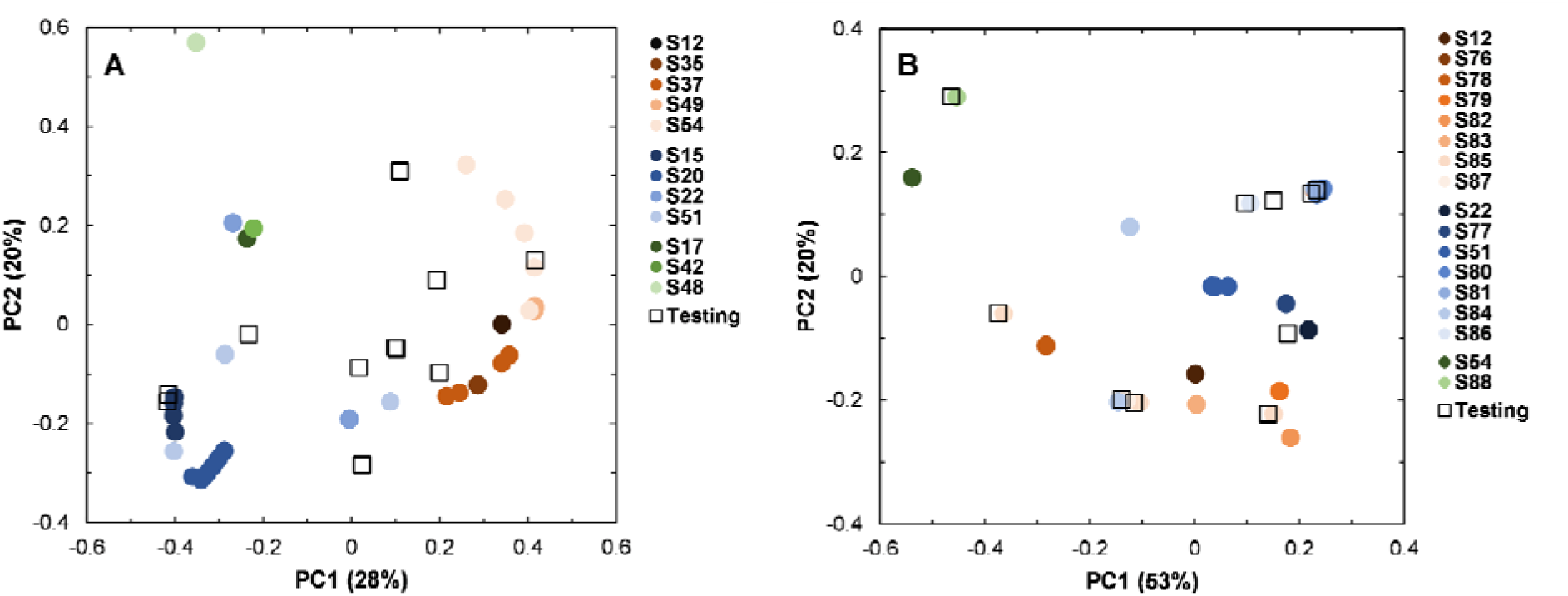
Bray Curtis distance-based PCoA of the training and testing datasets for calibrating (A) microbial kinetic parameters and (B) internal resistance.

### 3.2 Validating the data-driven component

The ANNs for calibrating kinetic parameters were trained with operation-related features as th inputs and seven microbial kinetic parameters as the outputs (SI Table S2). The outputs included the maximum specific growth rates and maximum substrate utilization rates of the fermentative, electroactive, and methanogenic populations, as well as the mediator yield of the electroactive population. Their values were derived using a previously established mechanistic model,^28^ which also served as the mechanistic component of the hybrid model. Some of the parameter values were estimated to be constant for multiple samples (e.g., 15 d^-1^ for the maximum substrate utilization rate of the electroactive population in S48) because they were the boundary values determined based on previous studies.^31, 83, 84^ After training, ten-fold cross-validation was performed. The relative RMSEs between the mechanistically derived and data-driven inferred parameter values ranged from 3.8% to 15.5% with an average of 9.4%, which was slightly lower than the 10.2% obtained with the Bayesian network in our previous work and significantly lower than the 26.2% obtained with a null model (SI Figure S1).^64^ In addition to the more accurate inferences, the applicability of the data-driven component is also improved as no experimental data (i.e., prior system outputs and microbial population abundances) are needed for inference.

The ANN for calibrating internal resistance was trained with the features related to system configuration (SI Table S3) as the inputs and internal resistance as the output. The values of the internal resistance retrieved from the literature ranged from 8.6 to 1345.0 Ω, which could again reflect the diverse configuration and operation of the selected bioelectrochemical systems. Nevertheless, the trained ANN was able to accurately capture the statistical connection between the configuration-related features and internal resistance, resulting in a low relative RMSE of 3.3% in cross-validation. In comparison, the relative RMSE obtained with the null model (trained using averaged input and output values) was 21.5% (SI Figure S1). Among the features, the electrode distance and the ion strength of the electrolyte were inferred to be critical factors that had a greater effect on the internal resistance. For example, the microbial desalination cell in T8 with a 10-cm electrode distance and 0.05 M salt concentration in the desalination chamber had internal resistance of over 600 Ω, while the microbial desalination cell in S54 only had 8 Ω due to a much shorter electrode distance and higher ion strength of solution (4 M) in the desalination chamber.

### 3.3 Predicting the system performance

#### 3.3.1 Internal resistance

By feeding the trained ANN with the configuration-related features from the testing dataset, internal resistance was accurately predicted over a wide range (Figure 3) as indicated by the high R² (0.98) and the low relative RMSE (3.9%) between the predicted and observed values. The smallest and largest observed internal resistance in the testing dataset was 8.6 to 758.0 Ω, and they were predicted to be 10.2 and 783.5 Ω. The majority of the internal resistance values were measured to be around 200 Ω, and some of the predictions within this range were slightly inaccurate. Most of these samples were retrieved from the same publication in which the systems were operated at an anode ion strength of around 0.02 M. Moreover, the systems were fed with complex and undefined substrates that were different from the rest of the samples in the training and testing datasets. Substrate complexity can lead to complicated microbial metabolism and consequently affect internal resistance.^85, 86^ The prediction for systems fed with low anode ion strength or complex substrates remained to be improved with more data.

**Figure 3.**
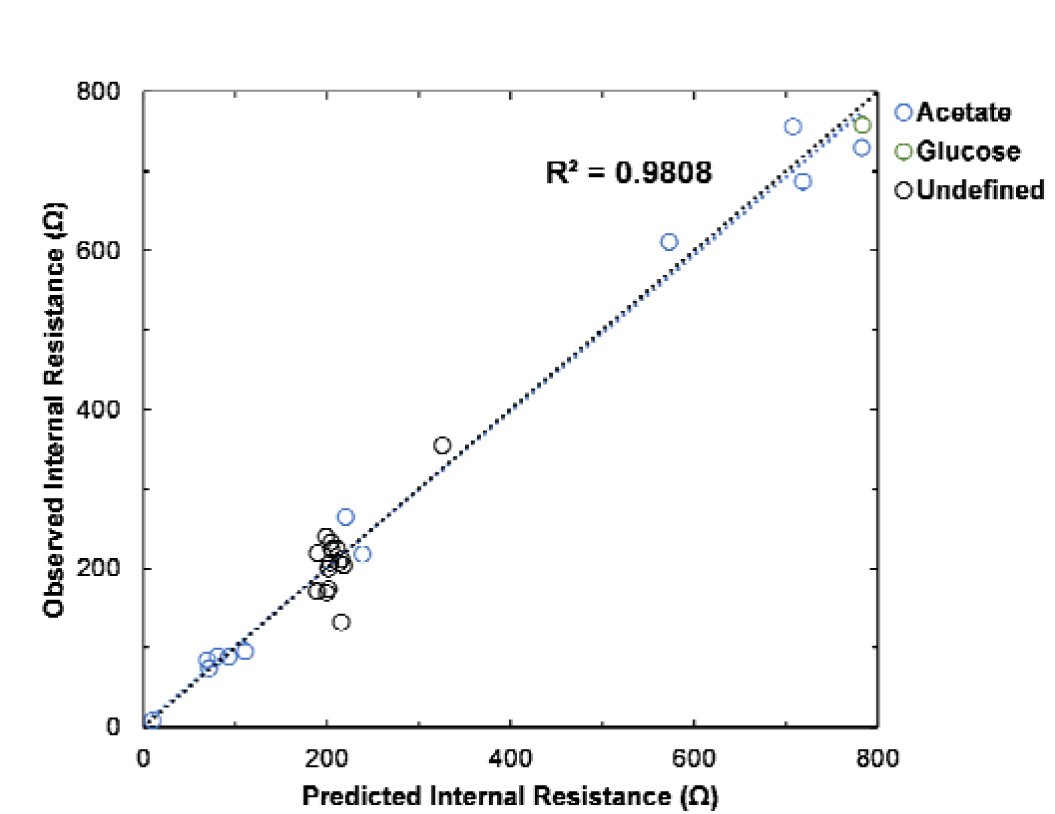
Comparison of the observed and predicted internal resistance.

Accurate prediction of internal resistance is highly desirable because it is a critical parameter that reflects the electron transfer kinetics.^74^ The internal resistance of bioelectrochemical systems can be divided into four main types: ohmic resistance of the system (e.g., electrode material and electrolyte conductivity), charge transfer resistance (i.e., the energy required to activate redox reactions), mass transfer resistance (e.g., caused by the diffusion of the substrates), and metabolism-related resistance (affected by the abundance and activity of the electroactive populations).^5^ It is challenging to model individual components mechanistically.^87^ While the metabolism-related resistance is often implicitly incorporated into multiplicative Monod expressions,^31, 87^ other types of internal resistance are lumped and parameterized in a statistical manner.^88, 89^ The parameters representing internal resistance are among the most influential parameters according to sensitivity analysis,^28^ and their calibration can significantly affect the mechanistic prediction of system performance. Data-driven approaches provide an experiment-free and reliable solution for calibrating internal resistance. The results suggest that the ANN can capture the complex relationship between system configuration and internal resistance, presenting a key step toward robust prediction of bioelectrochemical system performance.

#### 3.3.2 COD Removal

The seven microbial kinetic parameters (i.e., the maximum specific growth rates, maximum substrate utilization rates, and mediator yield) were calibrated by feeding the trained ANNs with operation-related features. The calibrated parameters and the internal resistance predicted in the previous section were then fed into the mechanistic component to simulate system performance (COD removal). The R² and relative RMSE between the observed and predicted COD removal were 0.81 and 7.3%, respectively (Figure 4). Two outliers were observed in the predictions. One of them was from a microbial electrolysis cell. In the training dataset, the number of samples for microbial electrolysis cells (38) was much less than that for microbial fuel cell samples (93), which might be the reason for the inaccurate prediction. The other outlier was from a publication (T1, SI Table S2) that studied the effect of electrode distance on the performance of microbial fuel cells. While the other five samples in the same study were accurately predicted (relative RMSE 2.6%), the cause of the outlier was not clear. Removing both outliers resulted in significant improvement in R² (0.90) and relative RMSE (5.2%) (Figure 4).

**Figure 4.**
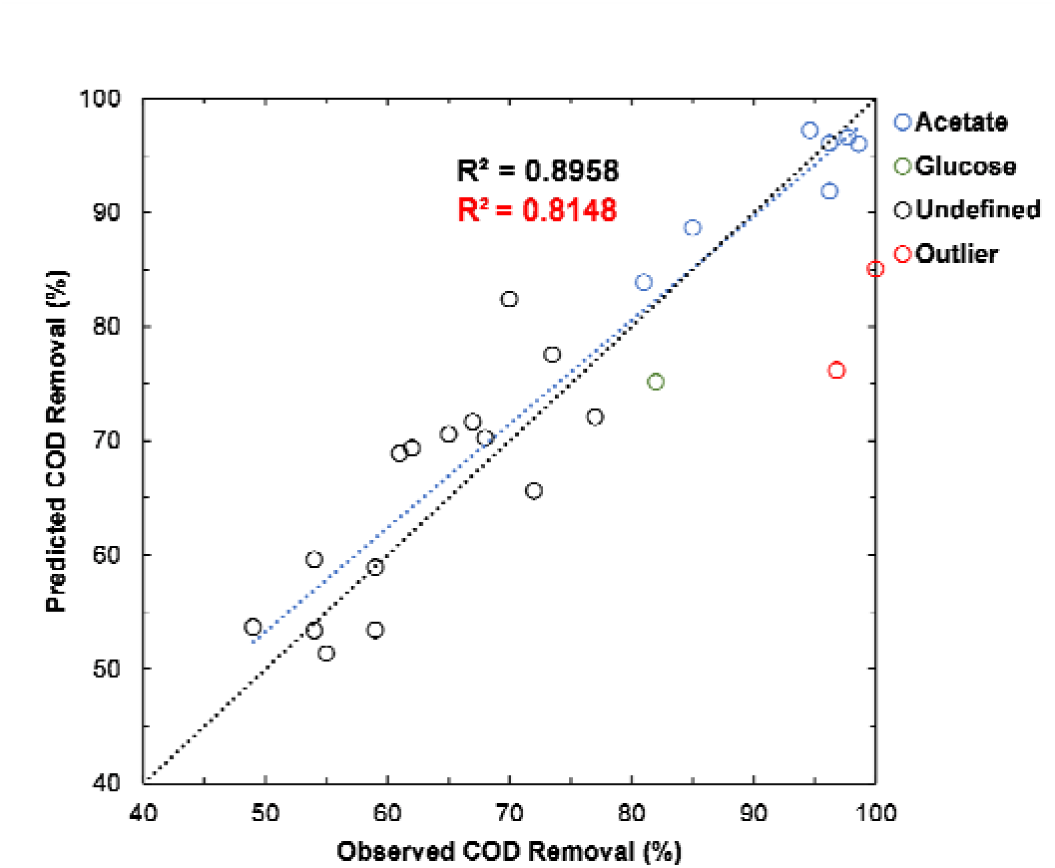
Comparison of the observed and predicted COD removal.

The prediction of COD removal was slightly less accurate than that of internal resistance because COD removal was predicted by coupling the data-driven and mechanistic components, during which the error could propagate.^90^ Nevertheless, the prediction performance of the hybrid model was much more robust than that of the control models. The standalone ANN yielded inaccurate predictions with an R^2^ of only 0.52 (SI Figure S2), and the standalone mechanistic model fed with averaged parameters could not yield any meaningful predictions (R^2^ 0.07, SI Figure S3). To further enhance the robustness of the hybrid model, more training data are needed, and each datum should contain a comprehensive set of features. Data availability from the literature is still lacking, calling for more standardized reporting and sharing of data.

#### 3.3.3 Current production

Four samples were selected from the testing dataset to predict real-time current production. Samples S5403 and T10_A were fed with acetate, T10_G was fed with glucose, and T2_1 wa fed with milk, representing three types of substrates (i.e., simple, fermentable, and complex) that were commonly used in the studies of bioelectrochemical systems. The current production of the acetate-fed systems was predicted with high R^2^ values (0.80 and 0.94) and low relative RMSE (4.5% and 12.7%) (Figures 5A & 5B). The prediction accuracy for the glucose-fed system dropped slightly with an R^2^ value of 0.77 and relative RMSE of 14.8% (Figure 5C). For milk-fed T2_1, the predicted current production did not agree with the observation, as indicated by the low R^2^ of 0.70 and high relative RMSE of 26.3% (Figure 5D). However, the total coulomb produced by milk-fed T2_1 was predicted with an acceptable error. This together with the well-predicted COD removal (Figure 5E) led to accurate prediction of coulombic efficiency (Figure 5F). The ability of the hybrid modeling strategy to capture indirect system performance such as total coulomb and coulombic efficiency warrants further investigation.

**Figure 5.**
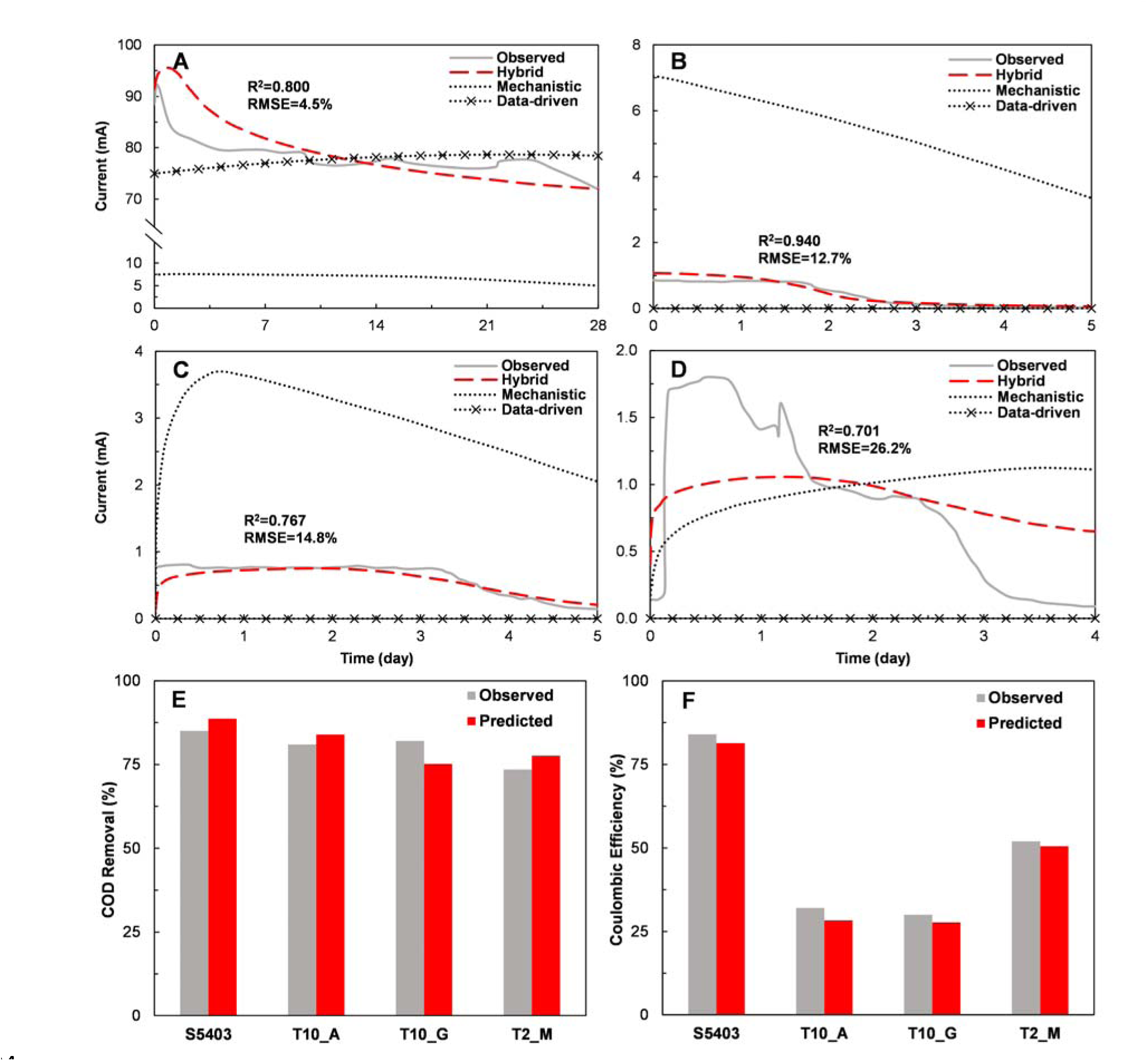
Observed and predicted current prediction of (A) sample S5403 fed with acetate, (B) sample T10_A fed with acetate, (C) sample T10_G fed with glucose, and (D) sample T2_M fed with milk. (E) COD removal and (F) coulombic efficiency predicted using the hybrid model.

The declined prediction performance with more complex substrates was likely a result of the variation in microbial metabolism and interactions.^85, 91^ For example, acetate has been shown to be a favorable substrate for electroactive populations.^92–94^ The resulting microbial communities are often dominated by these populations with a defined structure and show stable electroactivity. For systems that contain such communities, the statistical connection between environmental conditions and kinetic parameters can be better captured through data-driven modeling. Numerous studies have also shown how complex substrates act as deterministic factors to shape microbial communities.^95, 96^ The communities are dynamic in terms of structure and function, and the kinetic parameters are constantly changing, which can hardly be captured by a single prediction event. One of the potential solutions to improve the prediction accuracy is to frequently update microbial kinetic parameters based on real-time system outputs such as the effluent composition at different time points. However, such a dynamic prediction method requires a comprehensive training dataset containing high-quality timeseries data of all features. Despite the difficulty in modeling complex substrate-fed systems, the hybrid strategy noticeably outperformed standalone data-driven and mechanistic strategies. As shown in Figures 5A-5D, the control data-driven model and the mechanistic model fed with averaged kinetic parameters did not yield any meaningful predictions.

### 3.4 Perspectives of the hybrid modeling strategy

The ability to achieve experiment-free calibration of kinetic parameters suggests that the hybrid model can be used to design new bioelectrochemical systems *in silico*. To demonstrate this, a case study was conducted to optimize the configuration and operation of a single-chamber microbial fuel cell fed with 600 mg-COD/L acetate as the sole carbon source. While most of the features were held constant, five configuration-related features (anode/cathode electrode area, cathode electrode conductivity, electrolyte conductivity, and electrode distance) and one operation-related feature (hydraulic retention time) were set to different values (SI Table S4). These features were considered to be key variables because they differed significantly in the 169 publications reviewed in this study. Each configuration-related feature was set to up to eight values, and the resulting 11,120 combinations were fed into the ANN for predicting internal resistance. As shown in Figure 6, internal resistance could be reduced with a larger cathode area, a more conductive cathode electrode, a higher electrolyte conductivity, and a smaller electrode distance. Interestingly, an optimal anode area was predicted to be 1 m^2^. This is likely because a large anode electrode area can accommodate more electroactive biofilm, but when the area becomes too large, the resistivity of the electrode also increases.

**Figure 6.**
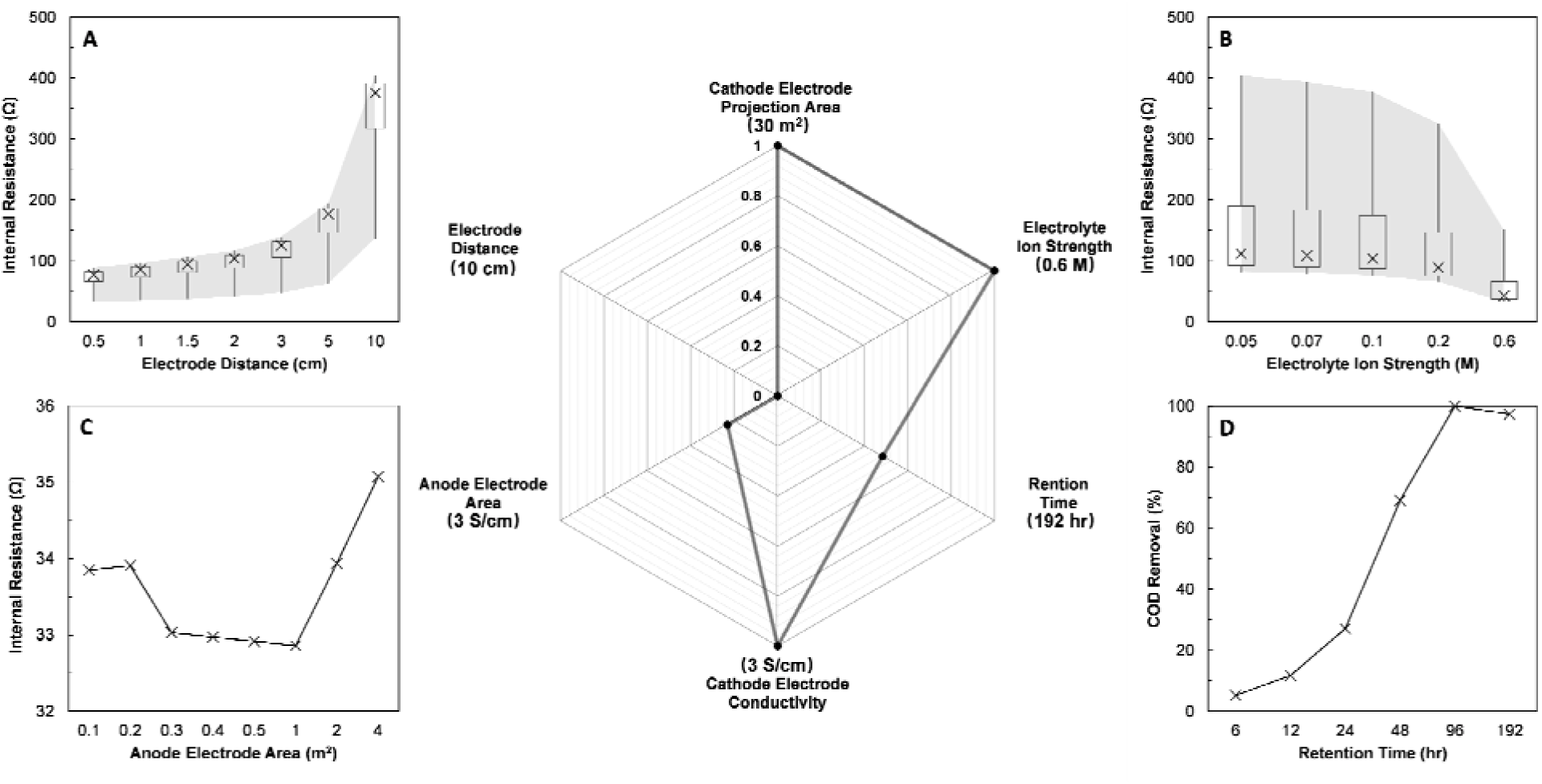
The internal resistance of a microbial fuel cell predicted with different (A) electrode distance, (B) electrolyte ion strength, and (C) anode electrode areas. (D) COD removal predicted with different hydraulic retention times. The numbers in the parentheses around the radar chart are the maximum values of the features. The numbers within the radar chart indicate the scale.

The minimum internal resistance of 32.9 Ω could be obtained with 0.5 cm electrode distance, 0.6 M electrolyte ion strength, 1 m^2^ anode electrode area (Figure 6A-C), 3 S/cm cathode electrode conductivity, and 30 cm^2^ cathode electrode area. This set of configuration-related features wa combined with retention time and fed into other ANNs to calibrate microbial kinetic parameters. The calibrated internal resistance and kinetic parameters were finally input into the mechanistic component to predict COD removal. As shown in Figure 6D, COD removal was predicted to increase from 5% to 69% when the retention time was prolonged from 6 h to 48h. COD could be completely removed with a retention time of 96 h. However, COD removal dropped slightly as the retention time was further increased to 192 h, possibly due to biomass decay and the release of soluble microbial products. Despite the great promise, the prediction performance of the hybrid model is still hindered by sample availability. Dynamic prediction of system performance is possible but requires a large training dataset with high-quality, comprehensive data for kinetic parameter estimation. The hybrid modeling strategy warrants further examination with different types of engineered biological systems such as full-scale activated sludge processes and anaerobic digesters.

## 4. CONCLUSIONS

In this study, a hybrid modeling strategy was developed to address the limitations of the previous strategies. The data-driven component in the hybrid model consisted of ANNs that were trained by computing errors with mechanistically derived parameter values. After training, the data-driven component took environmental conditions as the inputs to calibrate kinetic parameters, which were further fed into the mechanistic component for dynamic prediction of system performance. The strategy was examined using bioelectrochemical systems as a platform, and a hybrid model was built using 148 samples from 25 publications. When tested using 28 samples from seven additional publications, the hybrid model outperformed standalone data-driven and mechanistic models with accurate predictions of internal resistance, COD removal, and current production. The case study suggested that the hybrid model could potentially be used for *in silico* reactor design.

## Supporting information

Supporting_Infromation

Table_S5_References

## Acknowledgment

The authors would like to thank the “Sandwich” Doctoral Program at the Ben-Gurion University of the Negev, Israel, for supporting part of the research.

## GRAPHIC FOR TOC

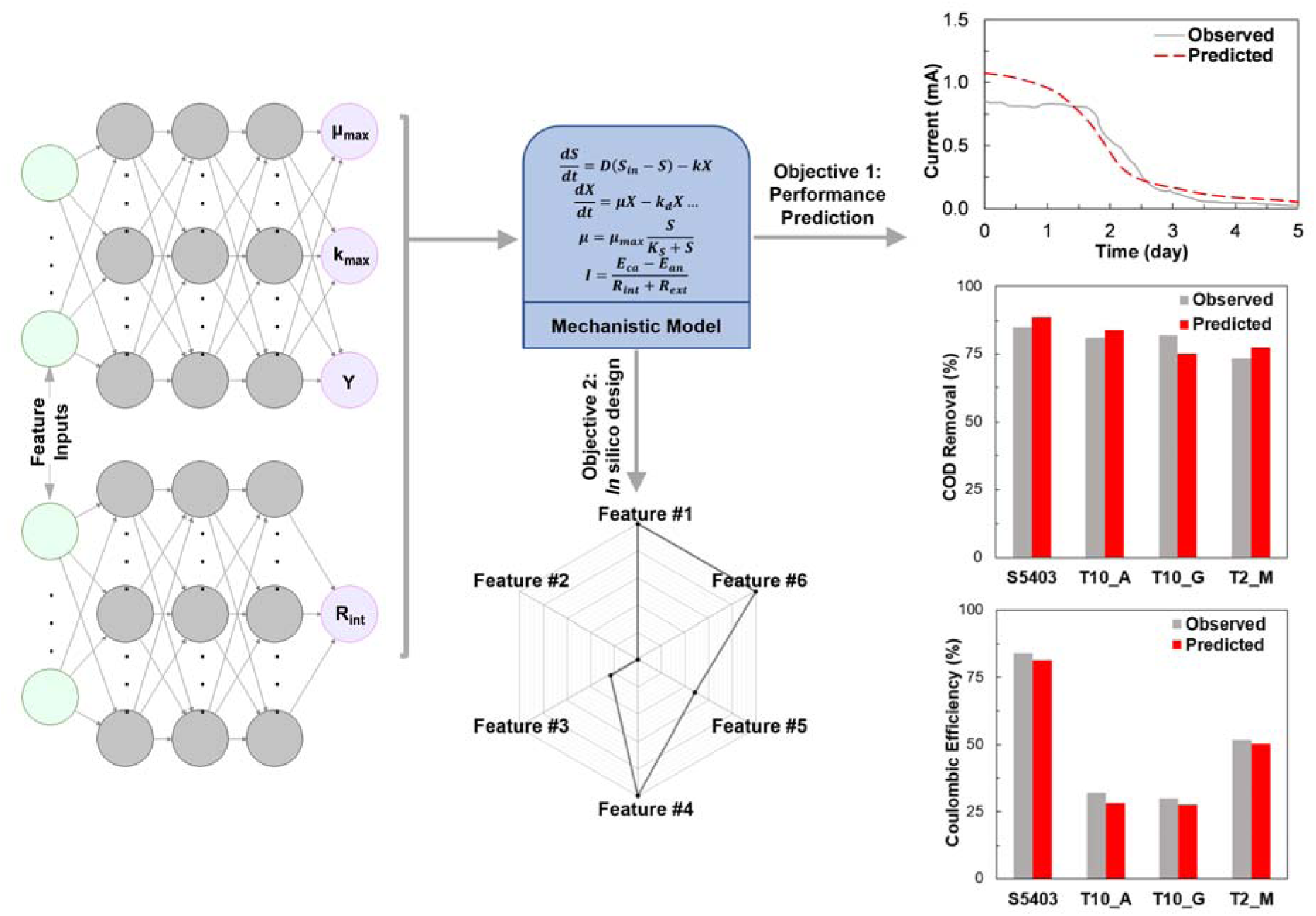

