## Supporting_Infromation for "Hybrid Modeling of Engineered Biological Systems through Coupling Data-Driven Calibration of Kinetic Parameters with Mechanistic Prediction of System Performance"

Intended for ***ACS ES&T Water***

Type of contribution***: Research Article***

^*^ Corresponding authors

**METHODS**

**Collection of training dataset**

We reviewed in total 169 publications and collected 148 samples from 25 publications for training the data-driven component [Supporting Information (SI) Table S1]. Because some of the samples lack key features to train the data-driven component for calibrating microbial kinetic parameters or internal resistance, the samples were divided into two groups to leverage the available data (some of the samples were used in both groups). Briefly, 73 samples from 12 publications were selected to train the ANNs for calibrating microbial kinetic parameters. The selection criteria were: 1) The publications reported operating conditions (e.g., substrate composition and concentration, pH, hydraulic retention time, etc.) and reactor configuration (e.g., anode area, external resistance, etc.). 2) The results included the variation of chemical oxygen demand (COD) over time or current/voltage that can be used to calculate the timeseries of COD. Meanwhile, 86 samples from 17 publications were selected to train the ANN for calibrating internal resistance. The selection criteria were: 1) The publications reported electrochemical conditions (e.g., anode area, ion strength, etc.) in addition to operating conditions and reactor configuration. Ion strength was estimated from Visual MINTEQ 3.1.^1^ 2) The results included internal resistance or measurements for calculating the internal resistance (e.g., polarization curves). Detailed information about the training dataset can be found in SI Tables S2 & S3. The samples in the training dataset are highly diverse in terms of system performance, operating conditions, reactor configuration, and electrochemical conditions, which is expected to enhance the applicability of the resulting model. To demonstrate the diversity of the samples, principal coordinate analysis (PCoA) based on Bray-Curtis distance was performed using the software R.

**Mass balance for primary degraders**

$$\frac{{dS}_{0}}{dt}=D\left( S_{0,in}-S_{0} \right)-k_{p}\frac{S_{0}}{K_{p}+S_{0}}X_{p} (Eq. S1)$$

$$\frac{{dX}_{p}}{dt}=\mu_{p}\frac{S_{0}}{K_{p}+S_{0}}X_{p}-k_{dp}X_{p}-D{\cdot X}_{p} (Eq. S2)$$

where $S_{0}$ is the concentration of complex substrates (mg/L); $S_{0,in}$ is the concentration of complex substrate in the influent (mg/L); $X_{p}$ is the concentration of the primary degraders (mg/L); $k_{p}$ is the maximum complex substrate utilization rate of the primary degraders (1/d);$\mu_{p}$ is the growth rate of the primary degraders (1/d); $K_{p}$ is the complex substrate half-saturation constant for the primary degraders (mg/L); $k_{dp}$ is the decay rate of the primary degraders (1/d); and $D$ is the dilution rate (1/d).

**Mass balance for electroactive and non-electroactive microbes**

$$\frac{{dS}_{1}}{dt}=D\left( S_{1,in}-S_{1} \right)+f_{e}\cdot k_{p}\frac{S_{0}}{K_{p}+S_{0}}X_{p}-k_{ee}\frac{S_{1}}{K_{ee}+S_{1}}\frac{M_{ox}}{K_{M}+M_{ox}}X_{ee}-k_{ne}\frac{S_{1}}{K_{ne}+S_{1}}X_{ne} (Eq. S3)$$

$$\frac{{dX}_{ee}}{dt}=\mu_{ee}\frac{S_{1}}{K_{ee}+S_{1}}\frac{M_{ox}}{K_{M}+M_{ox}}X_{ee}-k_{dee}X_{ee}-D\frac{1+\tanh\left( f_{ee}\left( X_{ee}+X_{ne}-X_{ee,max} \right) \right)}{2}X_{ee}(Eq. S4)$$

$$\frac{{dX}_{ne}}{dt}=\mu_{ne} \frac{S_{1}}{K_{ne}+S_{1}}X_{ne}-k_{dne}X_{ne}-D\frac{1+\tanh\left( f_{ne}\left( X_{ee}+X_{ne}-X_{ne,max} \right) \right)}{2}X_{ne} (Eq. S5)$$

$$\frac{{dM}_{ox}}{dt}={-Y}_{M}\cdot k_{ee}\frac{S_{1}}{K_{ee}+S_{1}}\frac{M_{ox}}{K_{M}+M_{ox}}+\frac{\gamma\cdot I}{V_{a}\cdot F\cdot X_{ee}n_{e}} (Eq. S6)$$

where $S_{1}$ is the concentration of simple substrates (mg/L); $S_{1,in}$ is the concentration of simple substrate in the influent (mg/L); $X_{ee}$, $X_{ne}$, and $M_{ox}$ are the concentrations of the electroactive microbes and non-electroactive microbes (mg/L), and mediator (dimensionless), respectively; $k_{ee}$ and $k_{ne}$ are the maximum substrate utilization rate of the electroactive microbes and non-electroactive microbes (1/d), respectively;$\mu_{ee}$ and $\mu_{ne}$ are the growth rate of the electroactive microbes and non-electroactive microbes (1/d), respectively; $K_{ee}$, $K_{ne}$, and $K_{M}$ are simple substrate half-saturation constants for the electroactive microbes, non-electroactive microbes (mg/L), and mediator half-saturation constant for the electroactive microbes (dimensionless), respectively; $k_{d,ee}$ and $k_{d,ne}$ are the decay rate of the electroactive microbes and non-electroactive microbes (1/d), $X_{ee,max}$ and $X_{ne,max}$ are the maximum capacity of electroactive microbes and non-electroactive microbes in the anode; $f_{e}$ is the fraction of complex degradation for energy, 0.78;^2^ $Y_{M}$ is the mediator yield for electroactive microbes; $\gamma$ is the molecular mass of mediator (mg/mol); $I$ is the current trough the circuit of BES (A); $V_{a}$ is the anode volume (L); $F$ stands for Faraday constant (A/d·mol); and $n_{e}$ represent the number of electrons transfer when a mediator is used by electroactive microbes.

**Kinetic parameter calculation**

The maximum substrate utilization rate and maximum growth rate are considered to be the most critical values for simulation of engineered bioprocesses,^2^ and mediator yield is a unique parameter for BES. Those kinetic parameters were estimated with specific limits according to previous studies while other parameters were retrieved from literature.^3, 4^ To estimate the parameters, the total substrate concentration over time and the initial values of the kinetic parameters were collected, calculated, or estimated from the selected papers. Because biomass and mediator concentrations were not measurable and unavailable in the literature, some assumptions are made: 1) primary degraders have an initial concentration of 100 mg/L in all reactors, 2) electroactive and non-electroactive microbe grow evenly on the anode surface at an average thickness of 60 μm,^5, 6^ 3) the maximum biofilm capacity in BES was assumed to be 600 mg/L,^7^ and 4) the electroactive microbe fraction is proportional to the CE. For microbial electrolysis cells whose CE was higher than 100% because of the applied voltage, the concentration of electroactive microbes was assumed to be 500 mg/L. Because the dimension of the anode electrode was not directly provided in some of the studies, the area was estimated based on the specific area and size of the electrode.^8^  Finally, the total substrate concentration over time with required constant values and initials was fed to the mechanistic component to calibrate the kinetic parameters.

**Internal resistance calculation**

A polarization curve can be divided into three regions that represent activation loss, ohmic loss and concentration loss. The ohmic loss region, which is typically the middle part of a polarization curve, commonly shows a linear relationship between the voltage and current. The internal resistance can thus be estimated from the slope of the ohmic loss region.

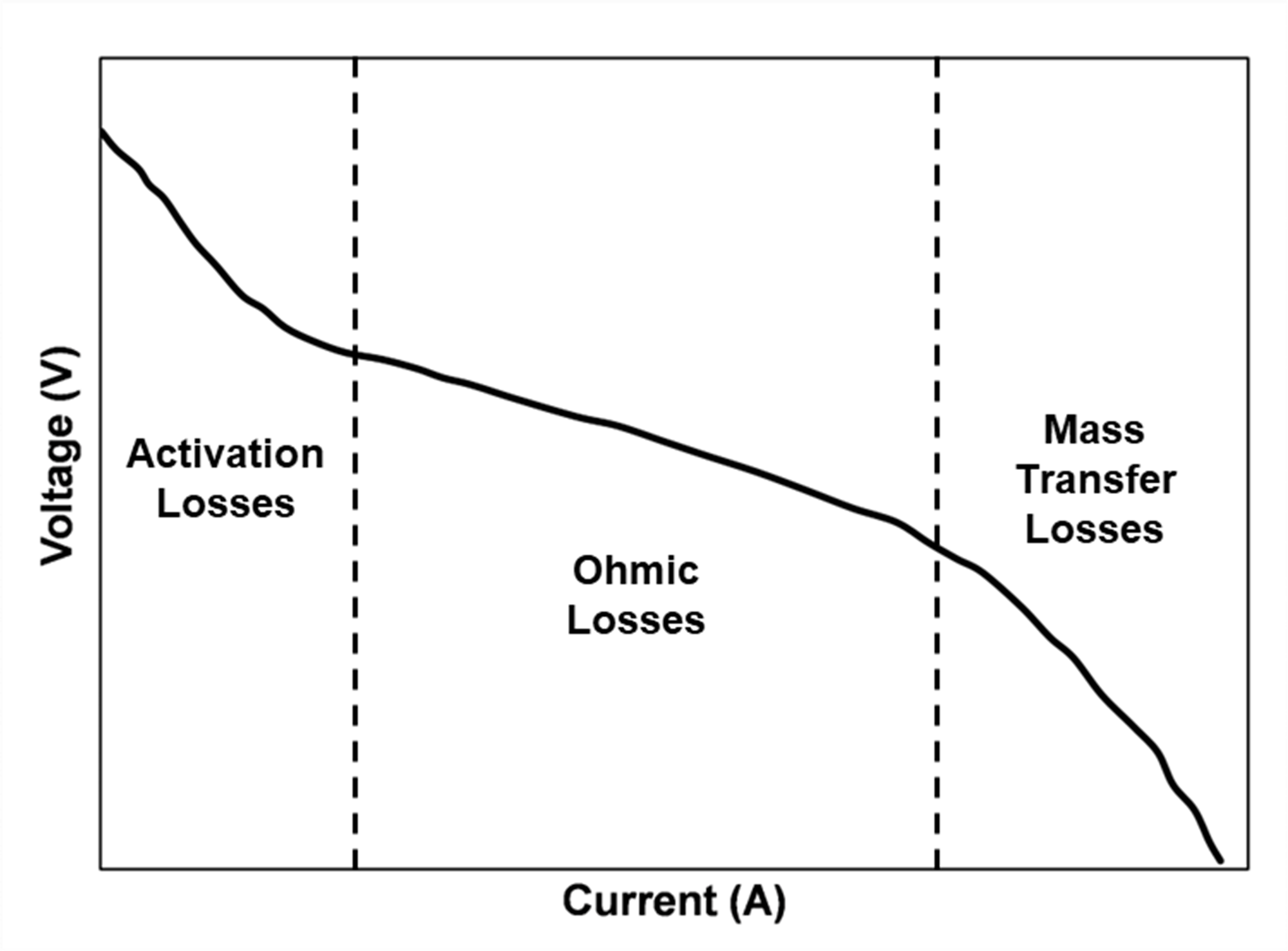

$$R_{int}=\left| \frac{V_{2}-V_{1}}{I_{2}-I_{1}} \right| (Eq. S7)$$

where *V_1_* and *V_2_* are the cell voltages at currents *I_1_* and *I_2_*, respectively, in the Ohmic loss zone of the polarization curve.

Figure S(M1). Polarization curve.

**Construction and validation of the data-driven component**

Before the data-driven component was trained, all the variable values were scaled to between 0 and 1:^9^

$$\hat{v}_{j,i}=\frac{v_{j,i}-v_{j,min}}{v_{j,max}-v_{j,min}} (Eq. S8)$$

where $\hat{v}_{j,i}$ is the normalized value of variable *j* in sample *i*, $v_{j,i}$ is the value of variable *j* in sample *i*, and $v_{j,max}$ and $v_{j,min}$ are the maximum and minimum values of variable *j*, respectively. Normalization is widely applied in data-driven modeling to make data of different scales comparable and to avoid bias.^10^ After normalization, the ANNs for calibrating microbial kinetic parameters were trained with operating conditions and reactor configuration as the inputs and mechanistically derived kinetic parameter values as the outputs (SI Table S2). The ANN for predicting internal resistance was trained with operating conditions, reactor configuration, and electrochemical conditions as the inputs and the calculated internal resistance as the output (SI Table S3). A time term was included as an additional training input based on the fact that the values of both microbial kinetic parameters and internal resistance could vary as a function of time. ANNs were trained using the R package Neuralnet with Sigmoid activation function.^11^ For simple training datasets, a three-layer ANN is sufficient to learn the statistical connection among the features. Empirically, the number of nodes in each hidden layer should be between the number of inputs and outputs.^12, 13^ Excessive hidden layers or nodes can lead to overfitting and compromise the prediction accuracy. Thus, the ANNs for calibrating microbial kinetic parameters and internal resistance both contained three hidden layers but eight and five nodes in each hidden layer, respectively.

Considering the dataset size and computational cost, ten-fold cross-validation was used to validate the ANN training method.^14, 15^ Briefly, the dataset was augmented with resampling to make its size an integer multiple of ten and was randomly split into ten subsets.^16^ For example, seven samples were first drawn from the 73-sample training dataset and then added back, making it in total 80 samples, which were then randomly divided into ten subsets with eight samples in each set. A validation ANN was trained with nine subsets, and the predictions were compared with experimental data in the remaining subset. Relative root-mean-square error (RMSE) and coefficient of determination (R^2^) were calculated as validation metrics.

$$relative RMSE=\frac{\sqrt{\frac{\sum_{1}^{n} \left( \hat{y}_{i}-y_{i} \right)^{2}}{n}}}{y_{max}} (Eq.S9)$$

where $\hat{y}_{i}$, *y_i_*, *y_max_* are the predicted, observed, and maximum observed values, respectively; and *n* is the number of samples. After validation, final ANNs were trained with the entire dataset. Null models were constructed as an additional validation step. This was performed by averaging all the inputs, kinetic parameters, and internal resistance across all samples.^17^

**Prediction of system performance**

To test the prediction performance of the hybrid model, 28 samples were collected from seven additional publications. Three types of system performance were predicted: internal resistance, COD removal, and current production. Internal resistance was predicted directly from the trained ANN and compared with the values derived from polarization curves. The majority of the input variables (*ENV* in Figure 1B) for predicting internal resistance were directly available in the publications. The time term (*∆t* in Figure 1B), on the other hand, had to be determined based on the operation duration before the polarization curves were measured. For example, if a polarization curve was measured immediately after a fresh substrate was added, *∆t* = 0. If a polarization curve was measured at the end of the operation cycle, *∆t* was then equal to the hydraulic retention time. Microbial kinetic parameters were inferred with the time term determined using the same method. Afterward, COD removal was calculated by inputting the calibrated internal resistance and microbial kinetic parameters into the mechanistic component. The initial values for mechanistic prediction were assigned according to the method reported in the previous study.^18^

Current production was predicted in a dynamic manner. To this end, the mechanistic component was modified by calculating current production using Ohm’s law:

$$I=\frac{E_{ca}-E_{an}}{R_{int}+R_{ext}} or I=\frac{E_{ap}}{R_{int}+R_{ext}}(Eq. S10)$$

Where $E_{ap}$ are the applied voltage to electrolysis cells, $E_{an}$ and $E_{ca}$ are respectively the anode and cathode potential of the other type of cells; $R_{int}$ and $R_{ext}$ are internal and external resistance, respectively. The anode and cathode potentials were calculated using the Nernst equation:

$$E_{an}=E_{ac}^{0}-\frac{RT}{8F}ln\frac{C_{{CH}_{3}{COO}^{-}}}{{C_{{HCO}_{3}^{-}}}^{2}\cdot\left( {10}^{-{pH}_{an}} \right)^{9}} (Eq. S11)$$

$$E_{ca}=E_{O_{2}}^{0}-\frac{RT}{4F}ln\frac{1}{p_{O_{2}}\cdot{{(10}^{-{pH}_{Ca}})}^{4}} (Eq. S12)$$

where $E_{ac}^{0}$ (0.187V vs. SHE) is the standard redox potential of acetate oxidation; $E_{O_{2}}^{0}$ (1.212 V vs. SHE) is the standard redox potential for oxygen reduction; *R* (8.314 J/K/mol) is the ideal gas constant; *F* (96485.3 A·s/mol) is the Faraday constant; T (K) is temperature; $C_{{CH}_{3}{COO}^{-}}$ and $C_{{HCO}_{3}^{-}}$ are molar concentrations of acetate and bicarbonate; $p_{O_{2}}$ is the oxygen partial pressure in the atmosphere. The pH in the anode and cathode was set to be constant by assuming an efficient buffer concentration. The anode potential was calculated based on the assumption that acetate was the main substrate for the electroactive populations to produce current.^2, 18-20^ To achieve dynamic prediction, an operation cycle was manually divided into several intervals, and the internal resistance within each interval was inferred by feeding the ANN with the corresponding *∆t*. Current production was therefore constantly updated within an operation cycle. Such dynamic prediction was carried out based on the fact that the substrate concentration changed with time, which could strongly impact the internal resistance. The prediction performance was tested by calculating the R^2^ and relative RMSE between the predicted and observed values. To further demonstrate the robustness of the hybrid model, standalone ANNs were trained with the same inputs but COD removal and current production as the outputs. Standalone mechanistic prediction was also performed by feeding the established mechanistic model with averaged microbial kinetic parameters and internal resistance of the 141 samples.

Table S1. Overview of the 25 and 6 publications for preparing the training and testing datasets, respectively.

| Study no. | Year | Reactor Type | Reference |
| --- | --- | --- | --- |
| S12 | 2016 | MDC | ^21^ |
| S15 | 2019 | Single-chamber MFC | ^22^ |
| S17 | 2017 | Two-chamber MFC | ^23^ |
| S20 | 2014 | Cassette MFC | ^24^ |
| S22 | 2015 | Cassette MFC | ^25^ |
| S35 | 2020 | Single-chamber MEC | ^26^ |
| S37 | 2019 | Single-chamber MEC | ^27^ |
| S42 | 2018 | Two-chamber MEC | ^28^ |
| S48 | 2021 | Two-chamber MEC | ^29^ |
| S49 | 2020 | Single-chamber MEC | ^30^ |
| S51 | 2019 | Single-chamber MFC | ^31^ |
| S54 | 2017 | MDC | ^32^ |
| S76 | 2017 | Single-chamber MFC | ^33^ |
| S77 | 2011 | Single-chamber MFC | ^34^ |
| S78 | 2017 | Single-chamber MFC | ^35^ |
| S79 | 2014 | Single-chamber MFC | ^36^ |
| S80 | 2020 | Single-chamber MFC | ^37^ |
| S81 | 2016 | Single-chamber MFC | ^38^ |
| S82 | 2022 | MDC | ^39^ |
| S83 | 2019 | Two-chamber MEC | ^40^ |
| S84 | 2021 | Single-chamber MEC | ^41^ |
| S85 | 2011 | Single-chamber MFC | ^42^ |
| S86 | 2015 | Single-chamber MFC | ^43^ |
| S87 | 2018 | Single-chamber MEC | ^44^ |
| S88 | 2018 | MDC | ^45^ |
| T1 | 2017 | Single-chamber MFC | ^46^ |
| T2 | 2015 | Single-chamber MFC | ^43^ |
| T4 | 2018 | Single-chamber MEC | ^44^ |
| T6 | 2011 | Single-chamber MFC | ^42^ |
| T8 | 2018 | MDC | ^45^ |
| T10 | 2011 | Single-chamber MFC | ^34^ |

Table S2. Features and microbial kinetic parameters of the 73 samples for training and 28 samples for testing (in the grey shadow).

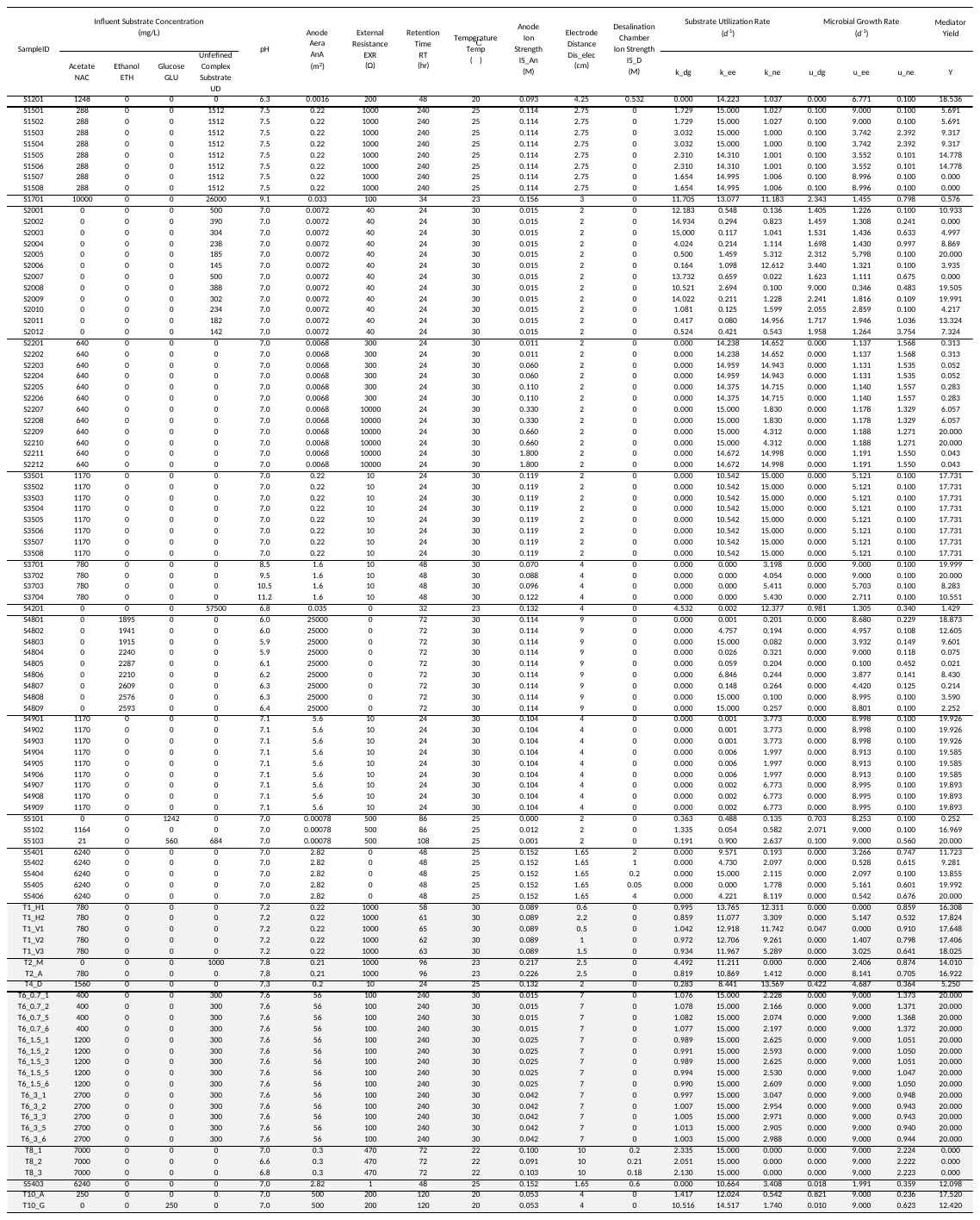
Table S3. Features and internal resistance of the 80 samples for training and 28 samples for testing (in the grey shadow).

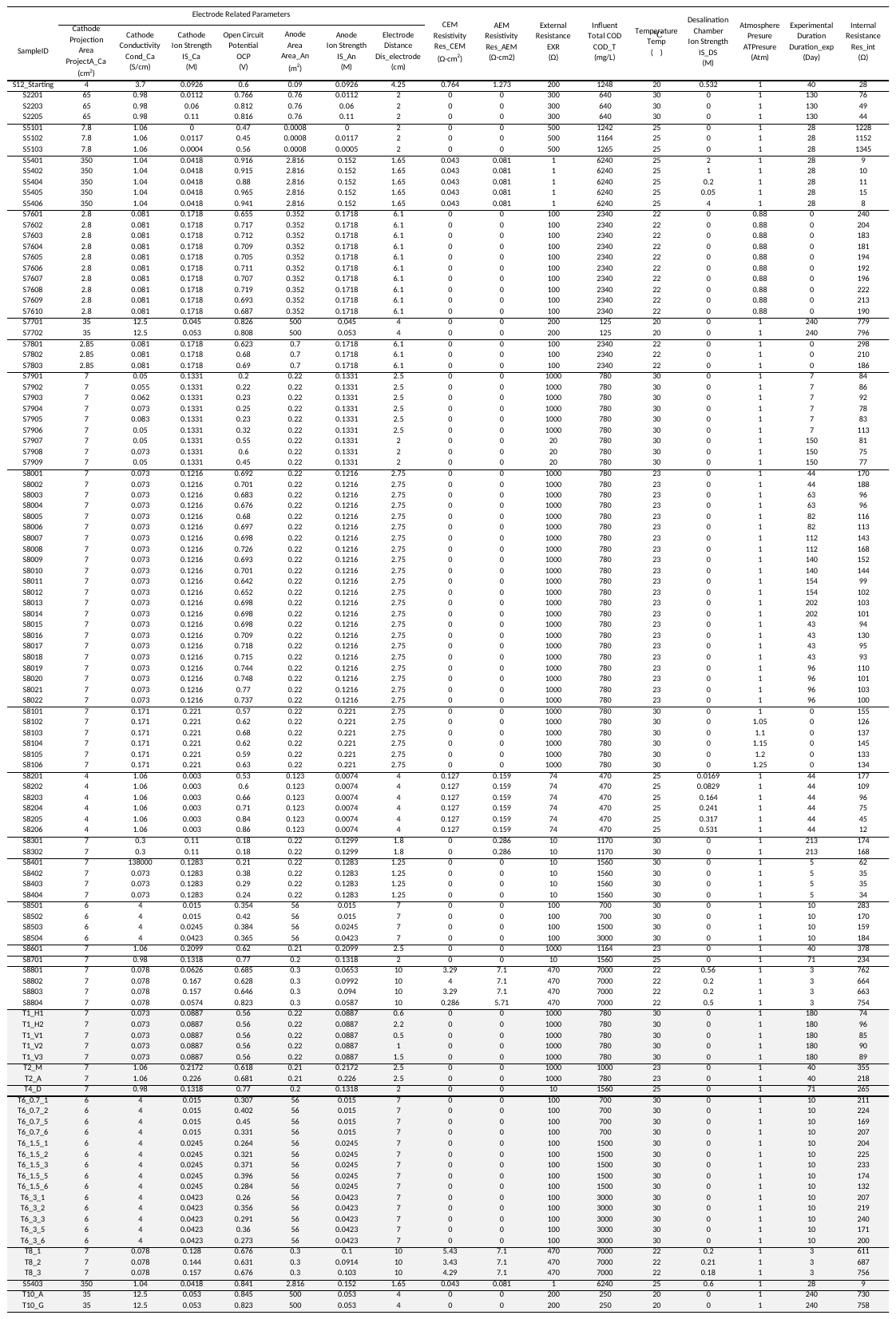

Table S4. Parameter values for the case study.

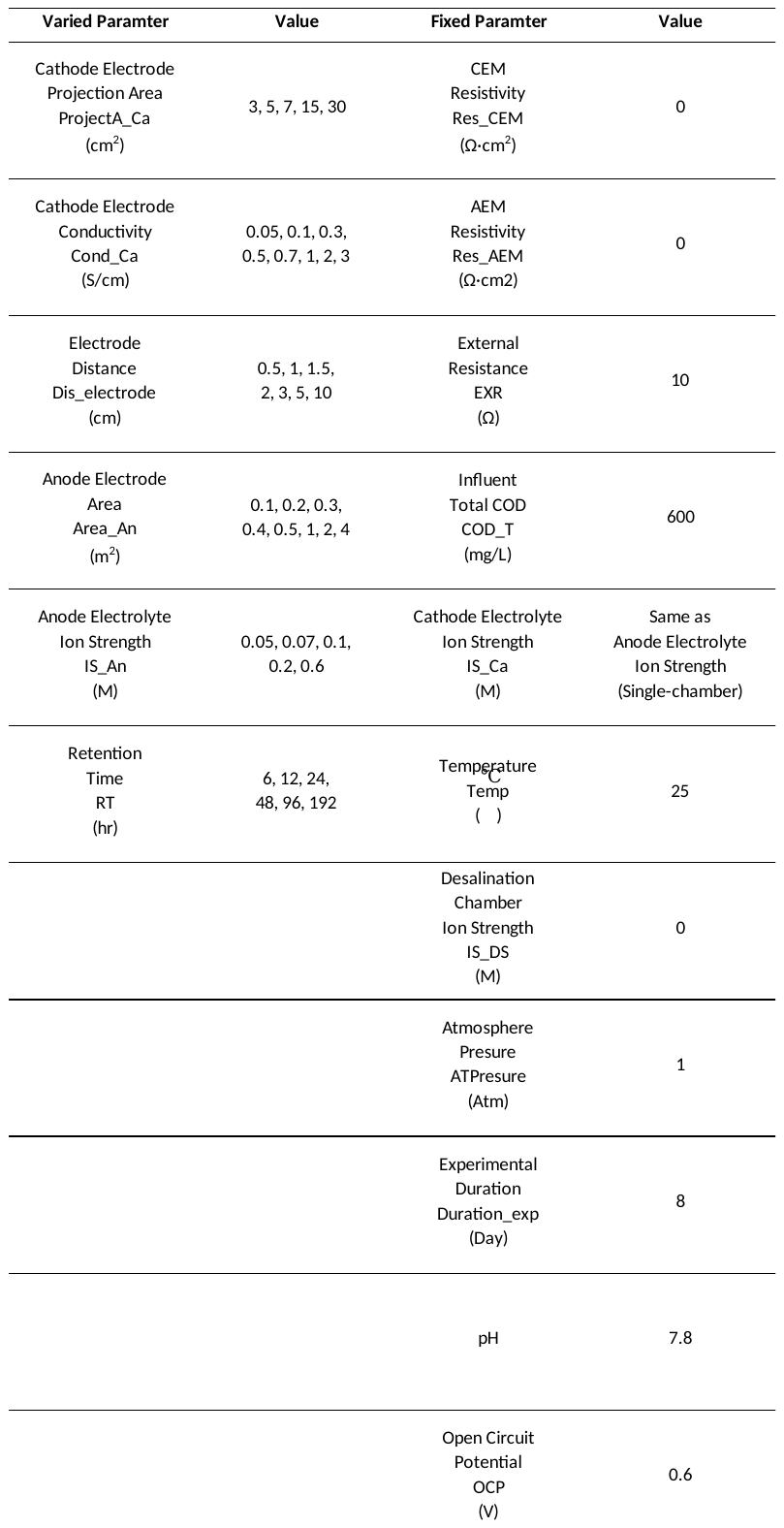

Table S5. R^2^ of the calibrated microbial kinetic parameters.

| **Sample ID** | **R^2^** |  | **Sample ID** | **R^2^** |  | **Sample ID** | **R^2^** |
| --- | --- | --- | --- | --- | --- | --- | --- |
| S1201 | 0.977 |  | S2203 | 0.912 |  | S4802 | 0.999 |
| S1501 | 0.985 |  | S2204 | 0.912 |  | S4803 | 0.968 |
| S1502 | 0.985 |  | S2205 | 0.912 |  | S4804 | 0.987 |
| S1503 | 0.941 |  | S2206 | 0.912 |  | S4805 | 0.995 |
| S1504 | 0.941 |  | S2207 | 0.901 |  | S4806 | 0.974 |
| S1505 | 0.962 |  | S2208 | 0.901 |  | S4807 | 0.989 |
| S1506 | 0.962 |  | S2209 | 0.869 |  | S4808 | 0.972 |
| S1507 | 0.879 |  | S2210 | 0.869 |  | S4809 | 0.962 |
| S1508 | 0.879 |  | S2211 | 0.921 |  | S4901 | 0.994 |
| S1701 | 0.974 |  | S2212 | 0.921 |  | S4902 | 0.994 |
| S2001 | 0.97 |  | S3501 | 0.933 |  | S4903 | 0.994 |
| S2002 | 0.988 |  | S3502 | 0.933 |  | S4904 | 0.867 |
| S2003 | 0.984 |  | S3503 | 0.933 |  | S4905 | 0.867 |
| S2004 | 0.975 |  | S3504 | 0.933 |  | S4906 | 0.867 |
| S2005 | 0.956 |  | S3505 | 0.933 |  | S4907 | 0.963 |
| S2006 | 0.958 |  | S3506 | 0.933 |  | S4908 | 0.963 |
| S2007 | 0.921 |  | S3507 | 0.933 |  | S4909 | 0.963 |
| S2008 | 0.91 |  | S3508 | 0.933 |  | S5101 | 0.989 |
| S2009 | 0.921 |  | S3701 | 0.966 |  | S5102 | 0.895 |
| S2010 | 0.933 |  | S3702 | 0.911 |  | S5103 | 0.964 |
| S2011 | 0.974 |  | S3703 | 0.935 |  | S5401 | 0.991 |
| S2012 | 0.951 |  | S3704 | 0.94 |  | S5402 | 0.849 |
| S2201 | 0.881 |  | S4201 | 0.903 |  | S5404 | 0.908 |
| S2202 | 0.881 |  | S4801 | 0.991 |  | S5405 | 0.912 |
|  |  |  |  |  |  | S5406 | 0.951 |

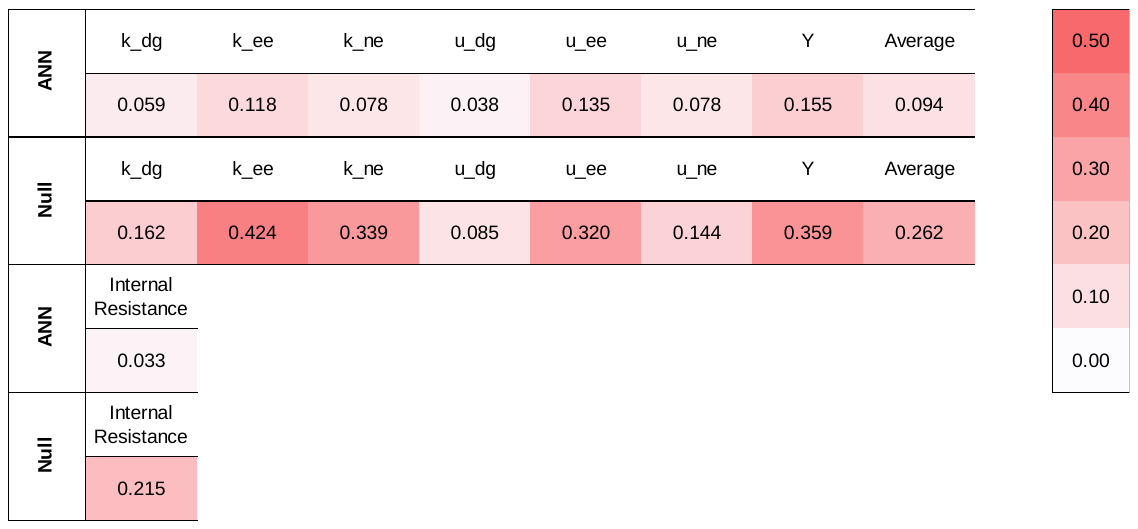

Figure S1. The relative RMSE of the kinetic parameters and internal resistance of the cross-validation.

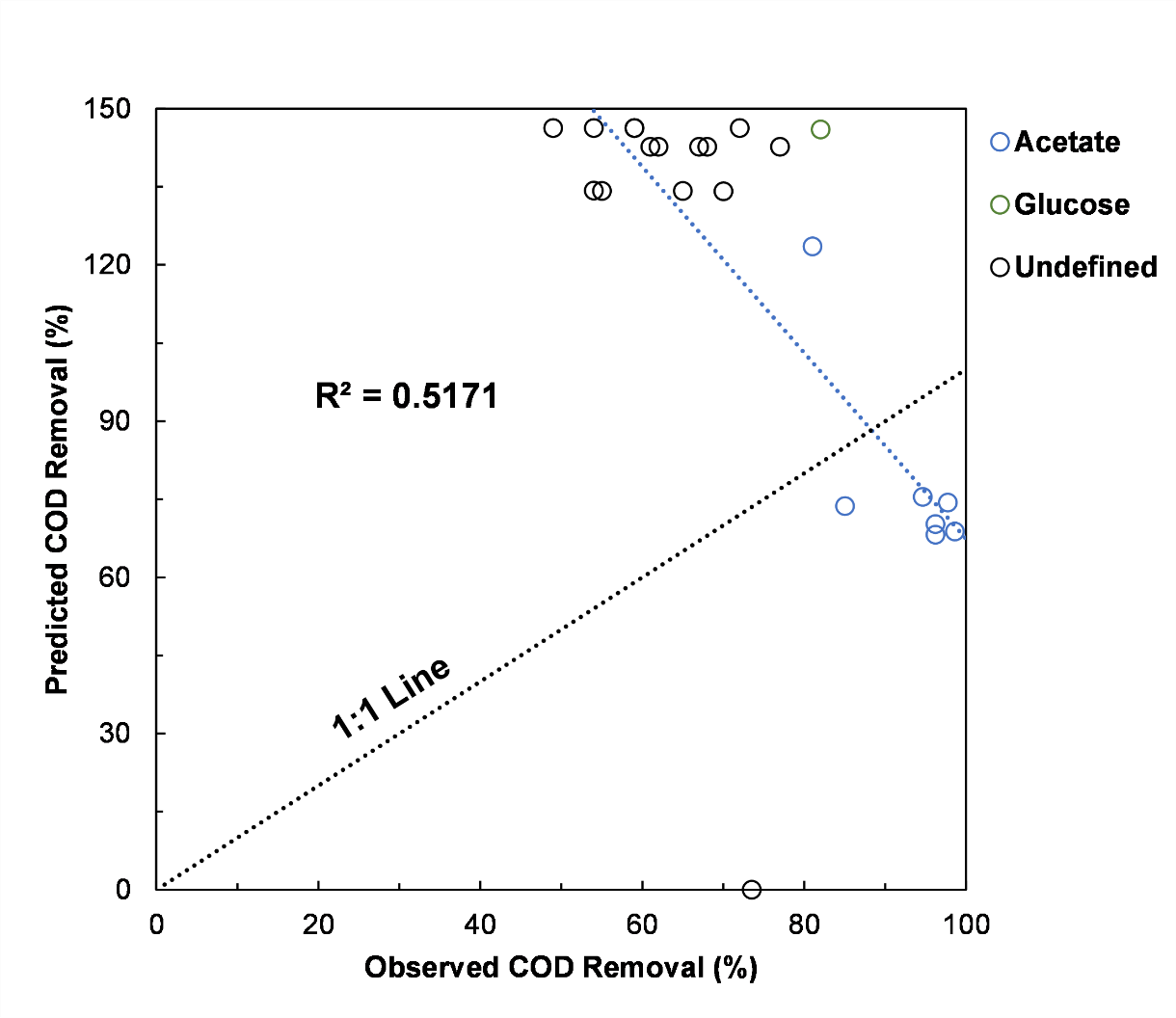

Figure S2. Comparison of the observed and predicted COD removal from pure data-driven model.

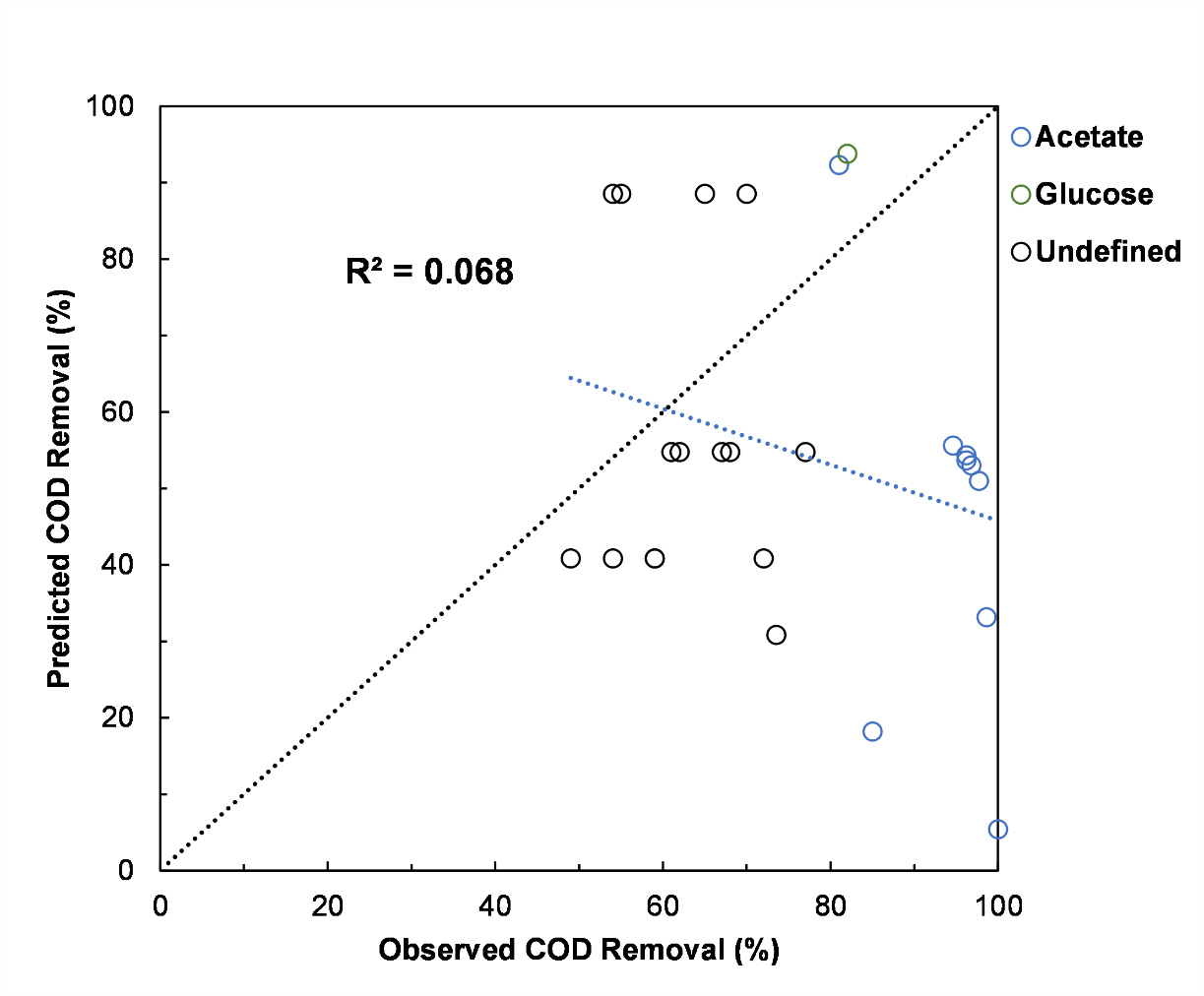

Figure S3. Comparison of the observed and predicted COD removal from mechanistic model with the inputs of kinetic parameter from the null model.
